# Personalized Molecular Signatures of Insulin Resistance and Type 2 Diabetes

**DOI:** 10.1101/2024.02.06.578994

**Authors:** Jeppe Kjærgaard Larsen, Ben Stocks, John Henderson, Daniel Andersson, Jesper Bäckdahl, Daniel Eriksson-Hogling, Jacob V. Stidsen, Kei Sakamoto, Kurt Højlund, Mikael Rydén, Juleen R. Zierath, Anna Krook, Atul S. Deshmukh

## Abstract

**Highlights:**

- Advanced proteomics analysis reveals personalized signatures of insulin resistance
- Fasting muscle proteome and phosphoproteome predicts whole-body insulin sensitivity
- Insulin-stimulated phosphoproteome reveals selective insulin resistance signatures
- Phosphoproteome and proteome atlas explains sex-specific muscle metabolism

Graphical Abstract

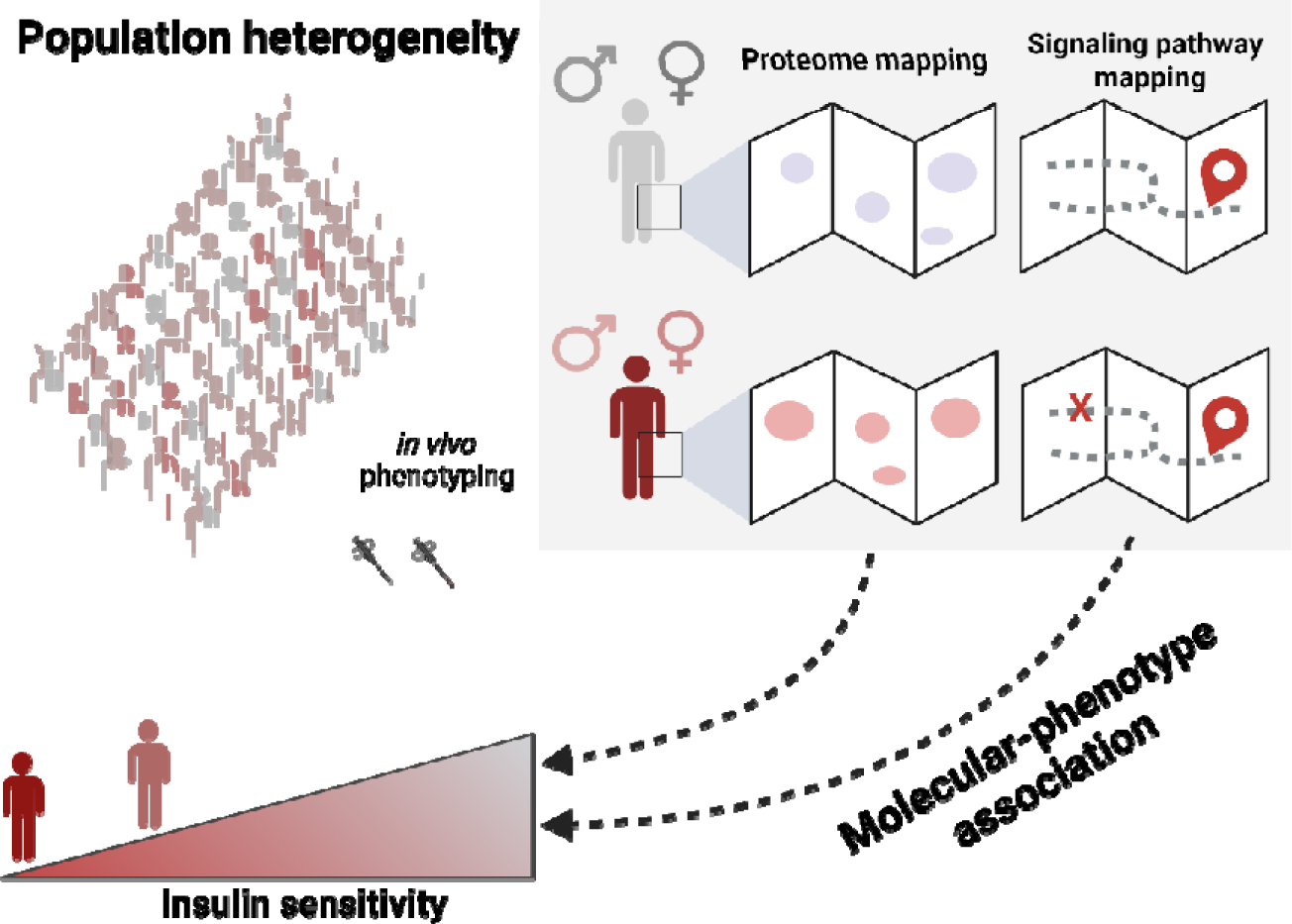

Insulin resistance is a hallmark of type 2 diabetes, which is a highly heterogeneous disease with diverse pathology. Understanding the molecular signatures of insulin resistance and its association with individual phenotypic traits is crucial for advancing precision medicine in type 2 diabetes. Utilizing cutting-edge proteomics technology, we mapped the proteome and phosphoproteome of skeletal muscle from >120 men and women with normal glucose tolerance or type 2 diabetes, with varying degrees of insulin sensitivity. Leveraging deep *in vivo* phenotyping, we reveal that fasting proteome and phosphoproteome signatures strongly predict insulin sensitivity. Furthermore, the insulin-stimulated phosphoproteome revealed both dysregulated and preserved signaling nodes - even in individuals with severe insulin resistance. While substantial sex-specific differences in the proteome and phosphoproteome were identified, molecular signatures of insulin resistance remained largely similar between men and women. These findings underscore the need for precision medicine approaches in type 2 diabetes care, acknowledging disease heterogeneity.

## Introduction

Type 2 diabetes is a growing global health challenge, with over 500 million cases worldwide (IDF Diabetes Atlas, 2023). Type 2 diabetes is characterized by elevated fasting or post-prandial blood glucose levels and peripheral insulin resistance, which affects liver, adipose tissue, and skeletal muscle. Thus, the pathogenesis of type 2 diabetes is remarkably heterogeneous, with disease progression influenced by both genetic and environmental factors^1^. Deep phenotyping and sub-group stratification have revealed that different clusters of type 2 diabetes are associated with clinical outcomes^2^. This complexity underscores the need for precision medicine approaches, in which individual differences are accounted for in the diagnosis, prevention, and treatment paradigms^1^.

Major advances have been made over the last decades to deduce the mechanism by which insulin regulates glucose uptake, as well as sites of resistance in type 2 diabetes^3^. Clinical studies utilizing the euglycemic-hyperinsulinemic clamp technique have established that skeletal muscle is quantitatively the main tissue involved in insulin-stimulated glucose uptake, and a major site of insulin resistance in type 2 diabetes^4–6^. Impaired insulin-stimulated glucose uptake has been attributed to a post-receptor defect, involving post-translational modifications and insufficient recruitment of GLUT4 to the plasma membrane^7,8^, rather than reduced abundance of signaling molecules or glucose transporters^9–11^. While targeted approaches have identified aberrant insulin signaling of selected molecules in skeletal muscle of people with type 2 diabetes^12–15^, a comprehensive system-wide investigation has yet to be performed. Furthermore, the extent to which individual variation in insulin signaling contributes to the heterogeneity of type 2 diabetes remains a knowledge gap.

Mass spectrometry-based proteomics has emerged as a powerful tool to elucidate mechanisms controlling cellular signaling and disease pathogenesis. This approach has been leveraged in cancer research and diagnostics, supporting the utility of proteomics and phosphoproteomics in precision medicine^16–18^. However, the lack of well-powered proteomics-focused studies of relevant tissues for insulin-resistance have hindered the implementation of precision medicine in type 2 diabetes. To understand the intricate heterogeneity within type 2 diabetes, a paradigm shift is required. By exploring variations in phenotypic traits, proteome and phosphoproteome signatures, and the responses to diverse environmental stimuli, alteration in causative proteins and pathways can be predicted, thereby enabling tailored approaches^19–22^.

Here we reveal features of skeletal muscle insulin resistance, previously unapparent using conventional comparative approaches. State-of-the-art proteomics technology and deep *in vivo* phenotyping were leveraged to map personalized diabetogenic traits with the skeletal muscle protein landscape in a cohort of >120 men and women with normal glucose tolerance or type 2 diabetes. We identify critical molecular pathways associated with insulin resistance. The molecular signature of skeletal muscle is strongly associated with clinical markers of insulin sensitivity, rather than fasting glucose control. Strikingly, the proteome and phosphoproteome landscape of skeletal muscle in the fasted state are critical determinants of whole-body insulin sensitivity. Furthermore, despite fundamental differences in substrate handling between men and women, the insulin resistant signatures are remarkably similar between sexes. These clinically focused data highlight the need to consider heterogeneity within disease classifications and advocate for a precision medicine approach to type 2 diabetes care.

## Results

### The proteome and phosphoproteome of human skeletal muscle are critical determinants of whole-body insulin sensitivity

To explore the molecular landscape of insulin resistance and type 2 diabetes, we recruited 77 participants (discovery cohort); consisting of 34 people living with type 2 diabetes (38% females) and 33 individuals with normal glucose tolerance (51% females). To validate our observations, we accessed samples from a published study^23^, of a further 46 participants (validation cohort); consisting of 34 people with type 2 diabetes (38% females) and 12 individuals with normal glucose tolerance (42% females) (Figure 1A). Clinical and physiological parameters for these cohorts are presented (Suppl. Table 1). Each cohort underwent deep *in vivo* glycemic-phenotyping using clinically relevant diagnostics, such as fasting glucose and HbA1c (measures of acute and chronic blood glucose control), as well as fasting insulin, HOMA1-IR (an estimation of insulin resistance derived from fasted insulin and glucose), and the hyperinsulinemic-euglycemic clamp derived M-value (a sensitive measure of insulin sensitivity). Skeletal muscle biopsies were collected both before and during the hyperinsulinemic-euglycemic clamp, allowing for the assessment of proteomic and phosphoproteomic molecular signatures within individuals in the fasted state, as well as the dynamics of acute insulin signaling.

**Figure 1.**
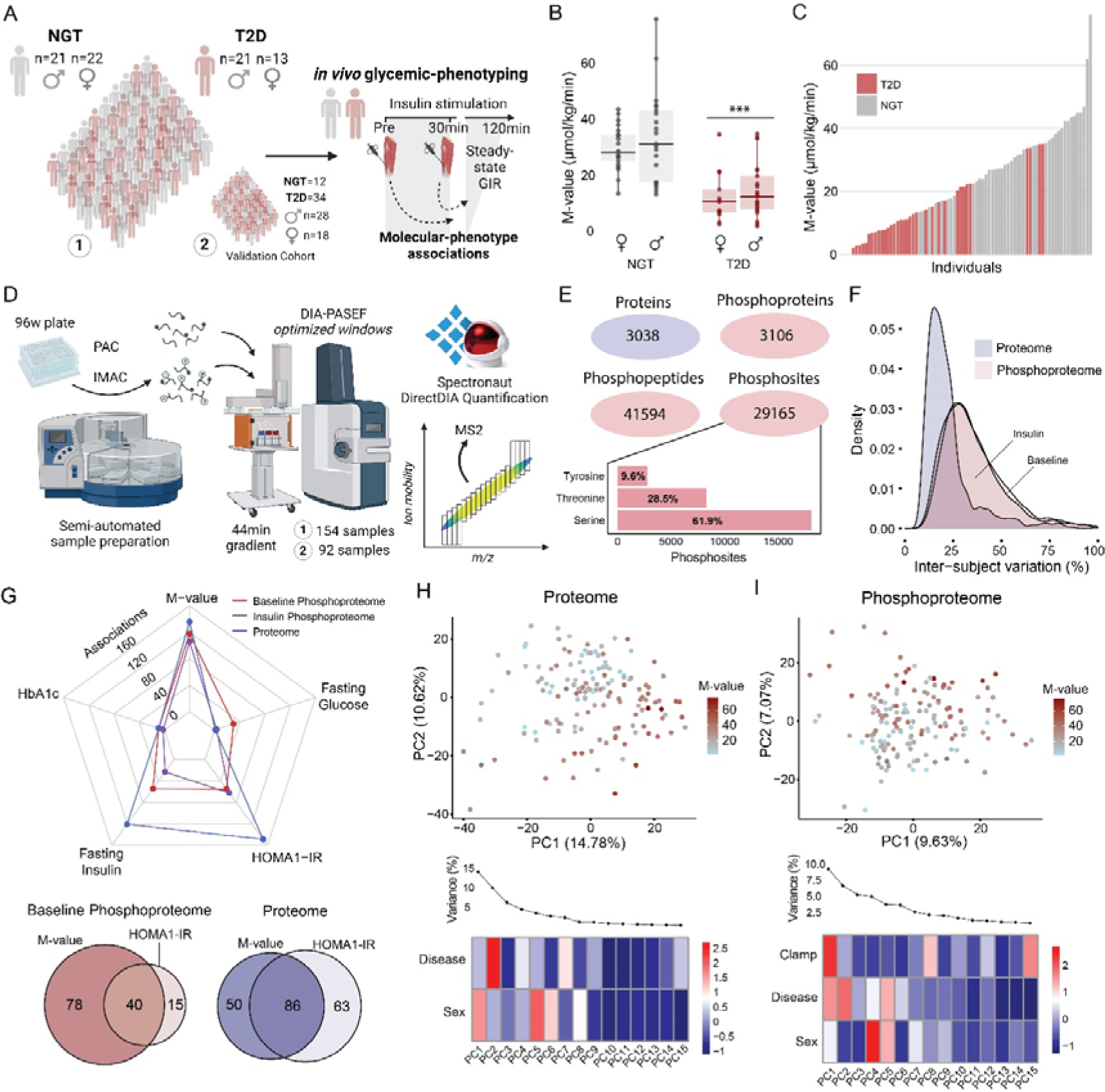
The Proteome and Phosphoproteome of Human Skeletal Muscle are Critical Determinants of Whole-Body Insulin Sensitivity. Schematic of study design: We recruited 77 individuals with normal glucose tolerance (NGT) (n=43; 21 male and 22 female) or type 2 diabetes (T2D) (n=34; 21 male and 13 female), and *vastus lateralis* muscle biopsies were collected pre and 30 minutes into an insulin clamp. A validation cohort of 46 individuals with NGT (n=12; 7 male and 5 female) or T2D (n=34; 21 male and 13 female) were also recruited (A). Boxplot of glucose-infusion rate during the steady-state period of the clamp, where the horizontal line indicates the median (B). A ranked bar plot demonstrating insulin sensitivity heterogeneity across all individuals (C). Reproducible and high-throughput (phospho)proteomics workflow on the KingFisher robot and evoseop-timsTOFpro liquid chromatography-tandem mass spectrometry setup. Samples were measured in DIA-PASEF mode and quantified in Spectronaut software (D). Number of proteins, phospho-proteins, peptides and sites quantified in at least 5 samples. Site phosphorylation distribution on serine, threonine and tyrosine residues (E). Inter-subject variation calculated as coefficient of variation across all individuals for proteome (baseline, fasting condition) and phosphoproteome (baseline and insulin) (F). Number of proteome (blue) and phosphoproteome (baseline = red, insulin = purple) associations with glycemic clinical measures. Venn diagrams depict overlap in associations between M-value and HOMA1-IR (G). Principal component analysis of proteome and phosphoproteome colored by M-value. Heatmap demonstrates z-scored PC loading contribution across disease, sex, and clamp (H-I). Insulin sensitivity associations were based on Kendall’s rank correlation with Benjamini-Hochberg corrected *P* values < 0.05 considered significant. GIR = Glucose infusion rate. NGT = Normal glucose tolerance. T2D = Type 2 diabetes. *P* < 0.001 = ***. All data presented in Figure 1 are from the discovery cohort.

Fasting glucose, HbA1C, fasting insulin and HOMA1-IR were elevated in people with type 2 diabetes (Figure S1A-B), while whole-body insulin sensitivity was reduced, as indicated by lower M-values (Figures 1B & S1B). Despite group differences, there was substantial heterogeneity in insulin sensitivity across the cohorts (Figures 1C & S1C), with M-values spanning a 36-fold difference in the discovery cohort (normal glucose tolerant individuals = 5.6-fold range, people with type 2 diabetes =16.7-fold range). Some individuals with type 2 diabetes even exhibited higher insulin sensitivity than some individuals with normal glucose tolerance (Figure 1C).

Intrigued by the observed heterogeneity in whole-body insulin sensitivity, we developed a high-throughput, semi-automated sample preparation workflow and analyzed the proteome and phosphoproteome of skeletal muscle using high-sensitivity mass spectrometry (Figure 1D). Leveraging recent advancements in short-gradient liquid chromatography coupled with the timsTOF pro2 in DIA-PASEF mode^24^, we efficiently measured the proteome and phosphoproteome from 77 individuals and 46 individuals in the discovery and validation cohort, respectively (a total of 492 mass spectrometry runs). This approach markedly advances capabilities beyond previous studies involving skeletal muscle phosphoproteomics. The phosphoproteomics screen covered 29,165 phosphosites (localization probability > 0.75, detected in at least 5 samples) on 3,000 phosphoproteins (Figure 1E). The proteomic analysis led to the quantification of 3,038 proteins (detected in at least 5 samples). After filtering for 25% valid values in each dataset, we quantified ∼3,000 proteins and ∼15,000 phosphosites within skeletal muscle, forming the basis for downstream bioinformatics analysis (Suppl. Table 2-3). The phosphoproteome and proteome coverage for the validation cohort (96 samples each) was comparable to the discovery cohort (Figure S1D, Supp. Table 3 & 4). The distribution of phosphosite residues revealed a prevalence of 62% for p-serine, 29% for p-threonine, and a noteworthy 10% for p-tyrosine. The substantial proportion of phospho-tyrosine (Y) residues emphasizes the unique phospho-tyrosine landscape for skeletal muscle compared to other tissues, which typically falls within the range of 0.1-2%^25,26^. The phosphoproteome exhibited greater inter-subject variability than the proteome (median = 35% vs median = 19%), suggesting a higher degree of heterogeneity at the phosphoproteome level (Figure 1F).

To evaluate the contribution of the skeletal muscle proteome and phosphoproteome to glucose homeostasis, we correlated the phosphoproteome (fasted, insulin-stimulated) and proteome (fasted) with the glycemic traits and clinical parameters (Figure 1G & Suppl. Table 6-8). Our findings revealed that the skeletal muscle proteome most strongly associates with insulin-dependent clinical measures, including fasting insulin, HOMA1-IR and M-value, whereas there was little to no association with fasting blood glucose and HbA1c. The phosphoproteome, both in the fasting state and during insulin-stimulation, exhibited more associations with the M-value than any other measures of insulin sensitivity, including HOMA1-IR. These data highlight the importance of skeletal muscle, and particularly phospho-signaling, in insulin-dependent glucose disposal. Furthermore, through visualization of the variance within the proteome and phosphoproteome, using a Principal Component Analysis (PCA), a continuum across different states of insulin sensitivity (M-value), rather than discrete grouping by diagnosis group, was observed (Figure 1H-I, Figure S1C-F). Categorical PCA loadings also highlighted diverse molecular drivers of separation, including biological sex. Thus, the human skeletal muscle proteome and phosphoproteome are linked to heterogeneity in whole-body insulin sensitivity, which may be dependent on sex.

### Proteomic Signature of Insulin Sensitivity in Human Skeletal Muscle

Capitalizing on the remarkable heterogeneity within the discovery cohort, we conducted a comprehensive analysis correlating protein levels with the M-value to elucidate the relationship between individual proteins and whole-body insulin sensitivity. This analysis yielded 136 associations (Kendall rank correlation, FDR < 5%) (Figure 2A & Suppl. Table 6), with HSPA2 and BDH1 displaying strong negative and positive correlations with insulin sensitivity, respectively (Figures S2A-B). We recapitulated our observations in the validation cohort with 89% conserved directionality of change (Figure S2C). Remarkably, only 15 proteins differed between people with type 2 diabetes or normal glucose tolerance in a discrete comparison (Figure S2D). This result highlights the pronounced variation in the proteomic landscape within the diagnosis groups and reinforces the need for precision diagnostics and therapeutics.

**Figure 2.**
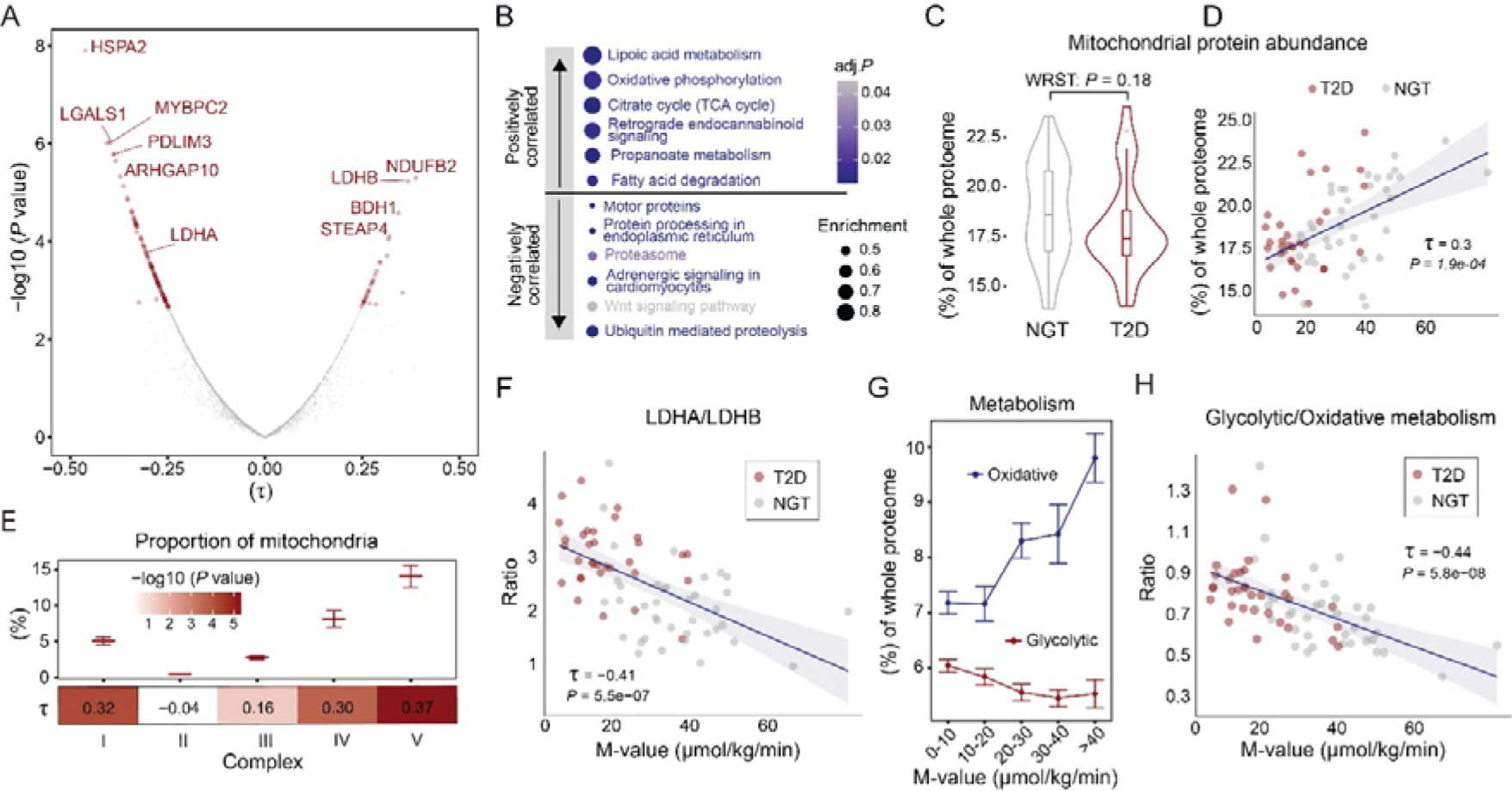
Proteomic Signature of Insulin Sensitive and Resistant Skeletal Muscle. Volcano plot of Kendall’s rank correlation coefficient (τ) of skeletal muscle proteome and insulin sensitivity associations (A). KEGG-pathway enrichment based on τ-ranked coefficients in positive and negative directions (adj.*P < 0.05*) (B). The proportion of summed mitochondrial protein abundance related to the whole proteome between individuals with NGT or T2D (C). Kendall’s rank correlation of mitochondrial abundance with insulin sensitivity (D). Proportion of electron transport chain complexes and association with insulin sensitivity adjusted for absolute mitochondrial protein (E). Association between the LDHA/LDHB ratio and insulin sensitivity (Kendall’s rank correlation) (F). The proportion of oxidative (Electron transport chain and TCA proteins) and glycolytic (glycolysis pathway proteins) metabolism proteins across a range of insulin sensitivity (G). The ratio of glycolytic/oxidative metabolism in skeletal muscle is inversely correlated with insulin sensitivity (Kendall’s rank correlation) (H). All data presented in Figure 2 are from the discovery cohort.

Proteins positively correlated with insulin sensitivity are linked to metabolic pathways such as oxidative phosphorylation and fatty acid degradation, commonly associated with insulin resistance and type 2 diabetes^27–29^. In contrast, the negatively correlated proteins are involved in processes that are generally underexplored in relation to insulin resistance, including proteasome and ubiquitin-mediated proteolysis, as well as Wnt- and adrenergic signaling. This suggests that altered protein degradation/turnover may contribute to the development of insulin resistance. Our investigation found no significant difference in mitochondrial protein abundance between the diagnosis groups (Figure 2C). However, mitochondrial protein abundance correlated with insulin sensitivity (Figure 2D), leading us to conclude that skeletal muscle mitochondrial abundance is not an inherent feature of type 2 diabetes, but rather related to insulin sensitivity. Notably, when adjusted for variations in mitochondrial abundance, the ATP-synthase complex (complex V) displayed the strongest correlation with insulin sensitivity among all the electron chain transport complexes (Figure 2E).

Lactate is an important metabolic intermediary in the transition between glycolytic and oxidative metabolism. The lactate dehydrogenase isoforms are functionally distinct and catalyze opposing reactions in the interconversion of lactate and pyruvate, with LDHA favoring the production of lactate (and therefore glycolytic metabolism) and LDHB favoring the production of pyruvate and entry of carbon intermediates into the TCA cycle for oxidative metabolism (Figure S2I). We found that the two isoforms of lactate dehydrogenase were regulated in opposing directions between the diagnosis groups, while LDH isoforms also had opposing associations with insulin sensitivity (Figure 2A & S2D-E). However, no difference was observed in the two major lactate transporters, monocarboxylate transporter 1 and 4, between groups (Figure S2F). The LDHA/LDHB proportion was strongly associated with insulin sensitivity (Figure 2F & S2G-H), which was more pronounced than the isoforms in isolation. This finding highlights the additional insight gained from examining stoichiometric relationships beyond the absolute abundance of individual proteins. To further dissect the metabolic machinery in skeletal muscle health, we calculated the total abundance for proteins involved in oxidative phosphorylation (TCA + electron transport chain) and glycolytic processes. We observed a clear increase in the abundance of oxidative phosphorylation proteins, from 7 - 10% comparing individuals with low to high insulin sensitivity (Figure 2G). Conversely, there was a reduction in the proportion of glycolytic proteins, indicating a shift in metabolic profile with increased insulin sensitivity. Moreover, we observed a negative correlation between the ratio of glycolytic/oxidative proteins and insulin sensitivity, indicating significant differences in the metabolic protein architecture across a spectrum of metabolic stages (Figure 2H).

### The Fasting Phosphoproteome Landscape is a Critical Determinant of Insulin Sensitivity

We next investigated the influence of the fasting/non-insulin-stimulated skeletal muscle phosphoproteome by performing a discrete comparison of skeletal muscle obtained from fasted individuals with type 2 diabetes or normal glucose tolerance. Our analysis revealed an alteration in 43 sites, including upregulation of VPS13B S206 and downregulation of ADD1 S358, both proteins studied in the context of type 2 diabetes and insulin action^30–32^ (Figure S3A). Kinase-enrichment analysis of the significantly altered phosphosites, revealed a downregulation of kinases associated with insulin signaling such as AKT, AMPK, RPS6KB1 (P70S6K), and RPS6KA2 (RSK-3) in individuals with type 2 diabetes. Conversely, kinases related to TGFβ and BMP signaling, including BMPR1A/B, CSNK2A2, TGFBR1, CAMK2G, and GSK3A, were upregulated (Figure S3B). Elevated TGFβ signaling has been linked to the development of type 2 diabetes in rodents and humans^33,34^.

Struck by the protein associations with insulin sensitivity in the fasting/basal state (Figure 2), we applied a similar approach to examine associations with protein-phosphorylation. We found 78 phosphosites correlating with insulin sensitivity exclusively in the fasted state, compared to 66 phosphosites exclusively in the insulin-stimulated state (Figure 3A & Suppl. Table 7). Additionally, 40 phosphosites sites correlated at both time points, indicating that these sites are already dysregulated at baseline, and therefore, the association with insulin sensitivity is independent of insulin stimulation (Figure 3A). Focusing on associations with insulin sensitivity, independent of insulin stimulation, we found that 93% of these associations displayed conservation of direction in the validation cohort (Figure S3C). These findings highlight the predictive power of the fasting phosphoproteome for whole-body insulin-stimulated glucose metabolism. As changes in total protein content can drive concurrent changes in post-translational modifications, we plotted phosphosite correlation coefficients alongside total protein quantification (Figure 3B). As expected, numerous phosphosites displayed concordant responses with the parent protein. However, a greater proportion of phosphosites were correlated with insulin sensitivity, even when the parent protein displayed no such correlation, indicating the importance of site occupancy in modulating the skeletal muscle signaling landscape. One example is GAPVD1 (also known as GAPEX-5) a known regulator of insulin-stimulated GLUT4 translocation in adipocytes^35^. Specifically, T390 on GAPVD1 was positively correlated with insulin sensitivity, while the parent protein did not correlate (Figure 3B). Identifying pathways and kinases associated with insulin sensitivity/insulin resistance is key in understanding the pathology of type 2 diabetes. Therefore, we performed a kinase-enrichment analysis on the phosphosites positively and negatively correlated with insulin sensitivity. While no specific kinases were linked to insulin sensitivity, the enrichment analysis revealed activation of JNK and p38 family kinases were associated with insulin resistance (Figure 4C). Although the JNK-p38 pathway has been implicated in inflammatory responses and the development of insulin resistance and type 2 diabetes^36–38^, our analysis identifies the JNK-p38 pathway as the main driver of aberrant skeletal muscle signaling in insulin resistance.

**Figure 3.**
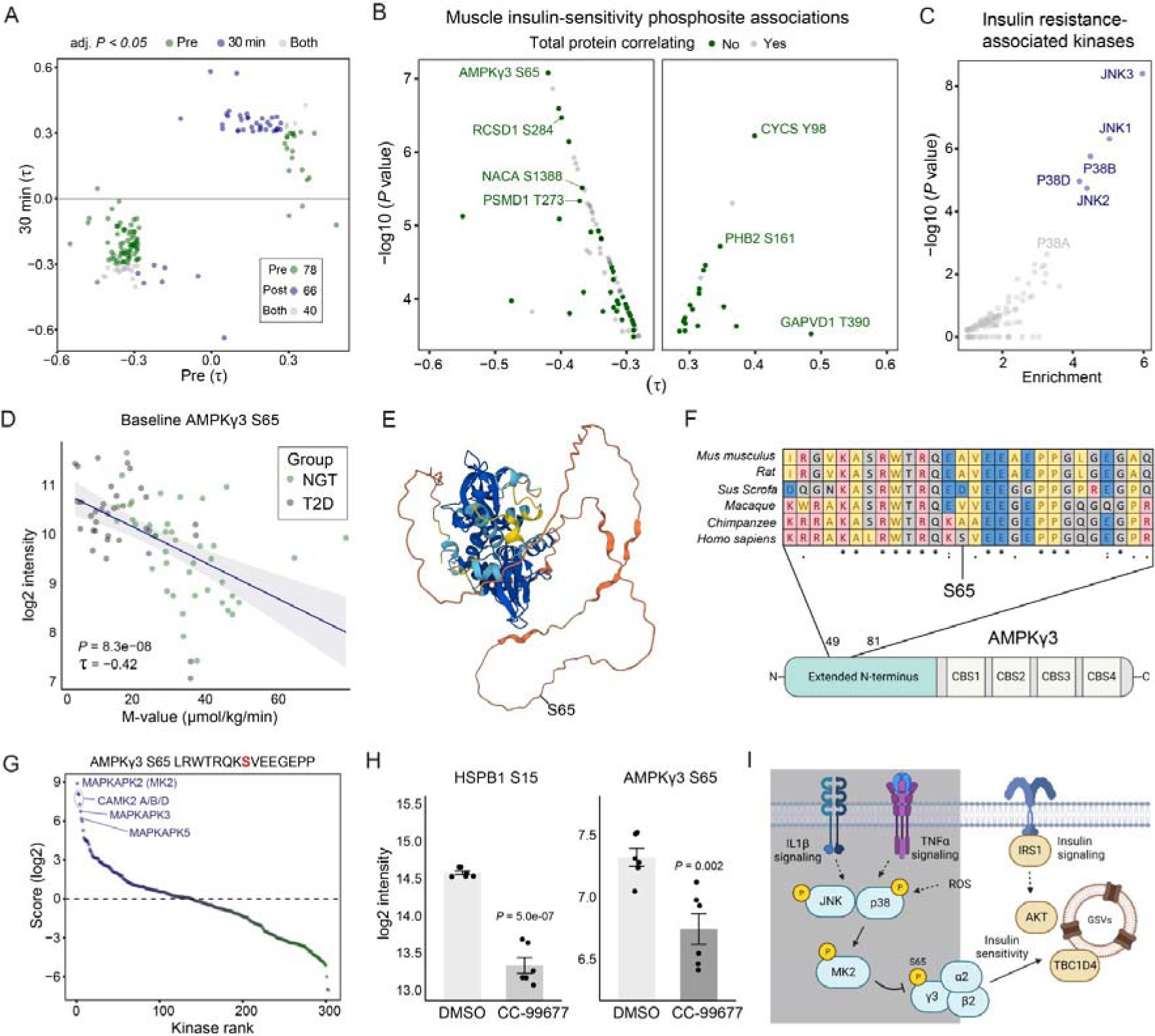
The Fasting Phosphoproteome Landscape is a Critical Determinant of Insulin Sensitivity. Phosphosite and insulin sensitivity associations at pre (fasting), 30 minutes into the insulin clamp (insulin-stimulated) and at both time points (A). Correlation coefficients of significantly correlated phosphosites with insulin sensitivity where the parent protein was quantified (B). The serine/threonine kinases that are predicted to be responsible for the phosphosites negatively correlated with insulin sensitivity in B (C). Baseline AMPKγ3 S65 phosphorylation with insulin sensitivity (D). Predicted alpha-fold structure of AMPKγ3 where the S65 residue is highlighted (E). Sequence alignment of AMPKγ3 residues 49-81 across species (F). Scoring of the ser/thr-kinome against the human AMPKγ3 S65 sequence (G). Log2 intensity of HSPB1 S15 and AMPKγ3 S65 phosphorylation from human skeletal muscle myotubes treated for 1 hour with DMSO or 1µM CC-99677 (H). Schematic of proposed signaling mechanism from inflammatory signals to AMPKγ3 S65 and to the regulation of skeletal muscle insulin sensitivity (I). The illustration was made in Biorender. MK2 = MAPKAPK2. All data presented in Figure 3 are from the discovery cohort.

**Figure 4.**
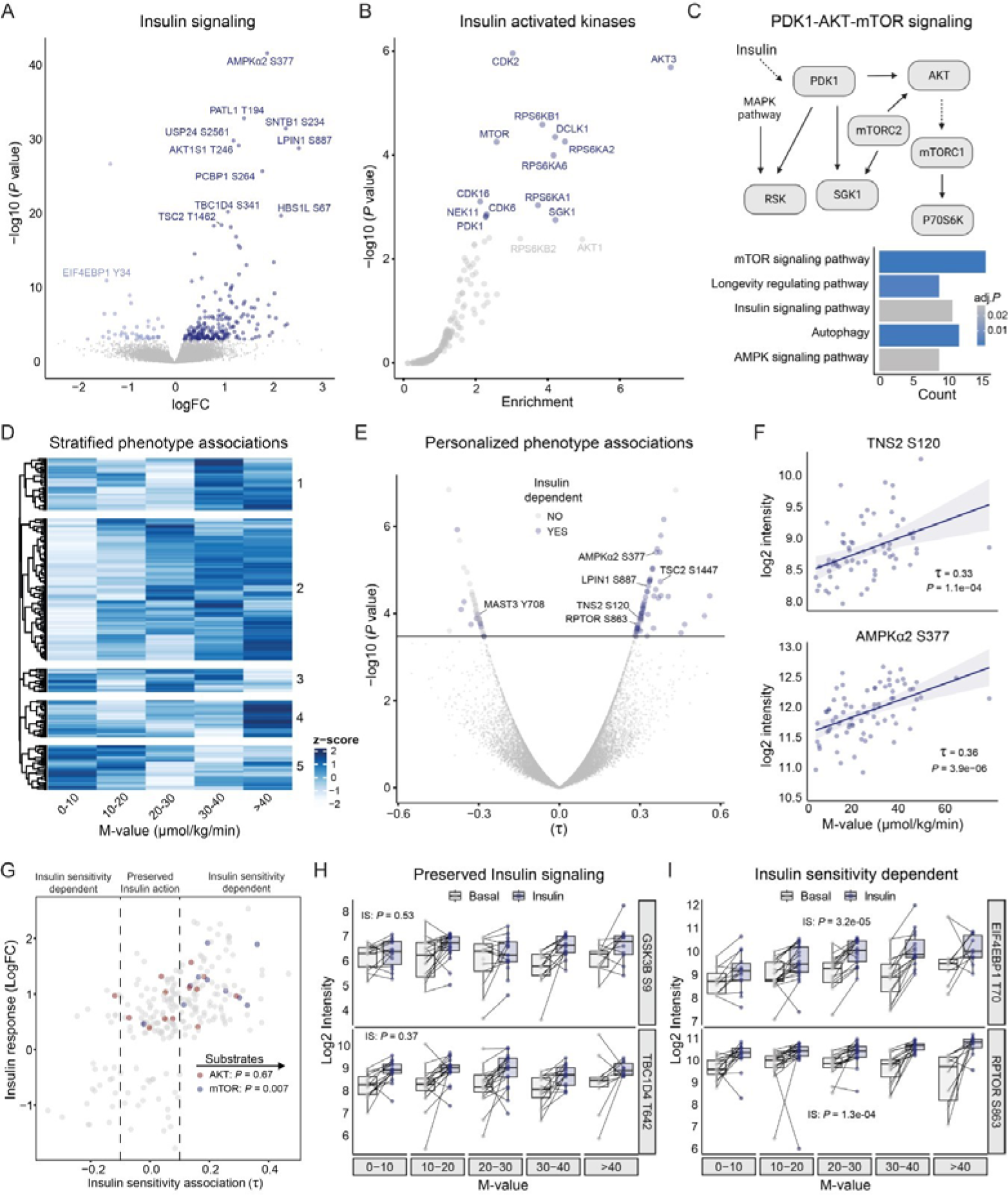
Preserved and Dysregulated Insulin Signaling Across States of Insulin Resistance. Volcano plot of insulin-regulated phosphosites (Limma main effect insulin, FDR <5%) (A). Unbiased prediction of skeletal muscle-expressed insulin-activated kinases. Enrichment was performed on all insulin-regulated phosphosites and whole unregulated phosphoproteome was used as background (B). Schematic network of insulin-activated kinases and overrepresentation enrichment analysis of signaling pathways (FDR < 5%) (C). Stratified insulin response across a range of insulin sensitivity. Unsupervised hierarchical clustering of insulin-regulated phosphosites was performed (D). Personalized insulin – phenotype associations. The 30 min phosphoproteome was correlated with individual insulin sensitivity (Kendalls rank correlation) (E). AMPKα2 S377 and TNS2 S120 phosphosite intensity association with insulin sensitivity (Kendalls rank correlation) (F). Insulin signaling-dependent and -independent associations with insulin sensitivity. Annotated AKT and mTOR sites are highlighted in red and blue, respectively. A Wilcoxon GeneSet test on insulin sensitivity correlation coefficient was performed to assess kinase enrichment (G). Examples of preserved AKT substrate phosphorylation (H) and insulin-sensitivity dependent mTOR substrate phosphorylation (I) across states of insulin sensitivity. All data presented in Figure 4 are from the discovery cohort.

The phosphosite with the strongest association with insulin resistance was S65 on the AMP-activated kinase (AMPK) regulatory γ-subunit 3 (Figure 3B & D). This finding was confirmed in the validation cohort (Figure S3 C-D). The S65 residue is located in the N-terminal tail of AMPKγ3, and this site is not present in the related γ1 and γ2 isoforms (Figure 3E). We verified the existence of this phosphorylation site through immunoprecipitation of N- or C-terminal FLAG-tagged human AMPKγ3 from HEK293 cell lysates, followed by subsequent proteomics analysis. (Figure S3E-F). Strikingly, when we compared the conservation of AMPKγ3 S65 across species, we found that this site was unique to *homo sapiens* and not present in the closest related species *Chimpanzee* (Figure 3F). However, in S*us Scrofa,* S65 is substituted with an aspartic acid (D), potentially serving as a biochemical or -physical mimic of constitutive phosphorylation. As AMPKγ3 is primarily a muscle-specific isoform, and the S65 is unique to humans, we further validated this phosphorylation event in primary human skeletal muscle myotubes. When we analyzed the human sequence surrounding AMPKγ3 S65, the top predicted kinases were MAPKAPK2 (MK2) and the CAMK2-family kinases (Figure 3G). We treated primary human skeletal muscle myotubes with the MAPKAPK2 inhibitor CC-99677 (Figure 3H). In addition to the expected decrease in HSPB1 S15 phosphorylation, a known substrate of MAPKAPK2, AMPKγ3 S65 phosphorylation was also reduced in response to CC-99677 treatment (Figure 4H).

MAPKAPK2 is a direct downstream target of p38^39^. Therefore, we propose that JNK-p38 pathway activation promotes insulin resistance via the AMPKα2β2γ3 trimer complex by indirectly phosphorylating S65 on the AMPKγ3 subunit through MAPKAPK2 (Figure 3I).

### Preserved and Dysregulated Insulin Signaling Across States of Insulin Resistance

Insulin signaling mediates a variety of metabolic and anabolic responses. Here we found 243 phosphosites to be regulated after 30 minutes of insulin stimulation (Figure 4A & Suppl. Table 8). These included increased phosphorylation of canonical insulin signaling proteins, such as TBC1D4 S341, AKT1S1 T246, and TSC2 T1462, and less-explored proteins like PATL1 T194. The insulin-regulated sites were enriched for regulatory functions related to protein activation, inhibition, localization, protein-protein interaction, and associations with disease (Figure S4A). An unbiased kinase enrichment analysis revealed activation of canonical insulin-activated kinases related to insulin and autophagy signaling, including PDK1, mTOR, AKT, SGK1, S6K (RPS6KB1), RSK (RPS6KA1, −3, −6) (FDR <5%, Figure 4B-C). In addition, several previously undescribed insulin-activated kinases were revealed, including DCLK1, CDK-2, −6, −16, and NEK11.

To extract deeper insight into phosphosite function, we associated insulin-stimulated phosphosite levels with insulin sensitivity. Using a stratified approach, we identified 5 clusters of phosphosites related to insulin sensitivity, the largest of which displayed a stepwise increase in insulin-stimulated phosphorylation with increasing insulin sensitivity (Figure 4D). At a personalized level, 66 phosphosites correlated with insulin sensitivity in response to insulin stimulation, with 54 and 12 sites positively and negatively correlated, respectively (Figure 4E). Notably, the phosphorylation of TNS2 (also known as TENC2 or C1-TEN) at S120 was positively correlated with insulin sensitivity only after insulin stimulation (Figure 4F & S4C). TNS2 is a tyrosine-phosphatase with a known role in counteracting INSR-induced IRS1 Y612 phosphorylation, muscle anabolic signaling, and GLUT4 regulation^40,41^. The positive correlation of AMPKα2 S377 (Figure 4E) and RPTOR S863, both mTOR substrate sites, suggests a role for mTOR in the insulin-dependent response to skeletal muscle glucose metabolism, consistent with a previous observation of skeletal muscle from young healthy men^22^. Furthermore, the phosphorylation of these sites in the fasting state did not correlate with insulin sensitivity, indicating that the dynamic response is specifically associated with insulin stimulation (Fig S4B-C).

Insulin-stimulated phosphosites displayed a large variability in the association to insulin sensitivity (Figure 4G). In fact, many sites with a large response to insulin were not associated with insulin sensitivity (−0.1 < τ < 0.1). These data indicate selective components of the insulin signaling pathway are preserved in states of insulin resistance. AKT and downstream substrates (e.g., TBC1D4) have been assayed as biomarkers of skeletal muscle insulin sensitivity. While phosphorylation of AKT2 S474, which is required for full activation of AKT, was associated with insulin sensitivity (Figure S4D), most of the *bona fide* AKT substrates displayed poor associations with insulin sensitivity (including GSK3B S9 and TBC1D4 T642) (Figure 4G-H). In contrast, mTOR substrates (e.g., EIF4EBP1 T70, RPTOR S863) were highly coupled to insulin sensitivity despite having diverse cellular functions in growth, translation, and autophagy^42–44^ (Figure 4G & I). These data highlight the need for a fundamental shift in the understanding of dysregulated signaling pathways during insulin resistance.

### The sex-specific molecular signature reveals shared and distinct features of metabolism

Hormonal and genetic factors influence skeletal muscle metabolism. Notably, the skeletal muscle transcriptome differs between males and females^45–47^, yet whether this is reflected at the protein level remains unclear. Herein, we illuminated the distinct molecular signatures of skeletal muscle from men versus women, providing a comprehensive sex-resolved atlas of the proteome and phosphoproteome (Suppl. Table 2 & 3). The women studied in this cohort were all post/peri menopausal, thus reducing possible cyclical effects of sex hormones. Principal component analysis showed a clear separation between males and females in the phosphoproteome and proteome (Figure S5A-B). Indeed, 110 proteins and 343 phosphosites were differentially regulated between sexes (Figure 5A-B). These major differences were recapitulated in our validation cohort, where 92% and 81% of the sex-specific proteins and phosphosites, respectively, shared the same directionality of change (Figure S5C-D). Differences in protein abundance included metabolically important proteins such as ALDH1A1 and DDAH1, as well as proteins targeted by FDA-approved drugs (Figure 5A & S5E). Systematic differences in substrate handling enzymes were also observed. Proteins related to glucose metabolism and oxidative phosphorylation were enriched in males, while proteins related to lipid uptake/storage were higher in females, consistent with increased plasma free fatty acids (FFAs) (Figure 5C & S S5F-G). The sex-associated phosphosites were located on signaling proteins with known site-regulatory functions in a multitude of biological processes. An example of this is the phosphorylation of S57 on Polyubiquitin-B (UBB), which was markedly higher in females. Phosphorylation of UBB S57 is a key event accelerating PARKIN-dependent mitophagy, suggesting a higher mitochondrial turnover in females^48,49^ (Figure S5H). Moreover, global analysis of the phosphoproteome revealed sex-specific enrichment for kinases, suggesting distinct activation of signaling (Figure 5D). These results highlight the importance of considering sex as a biological variable in study design and clinical trials.

**Figure 5.**
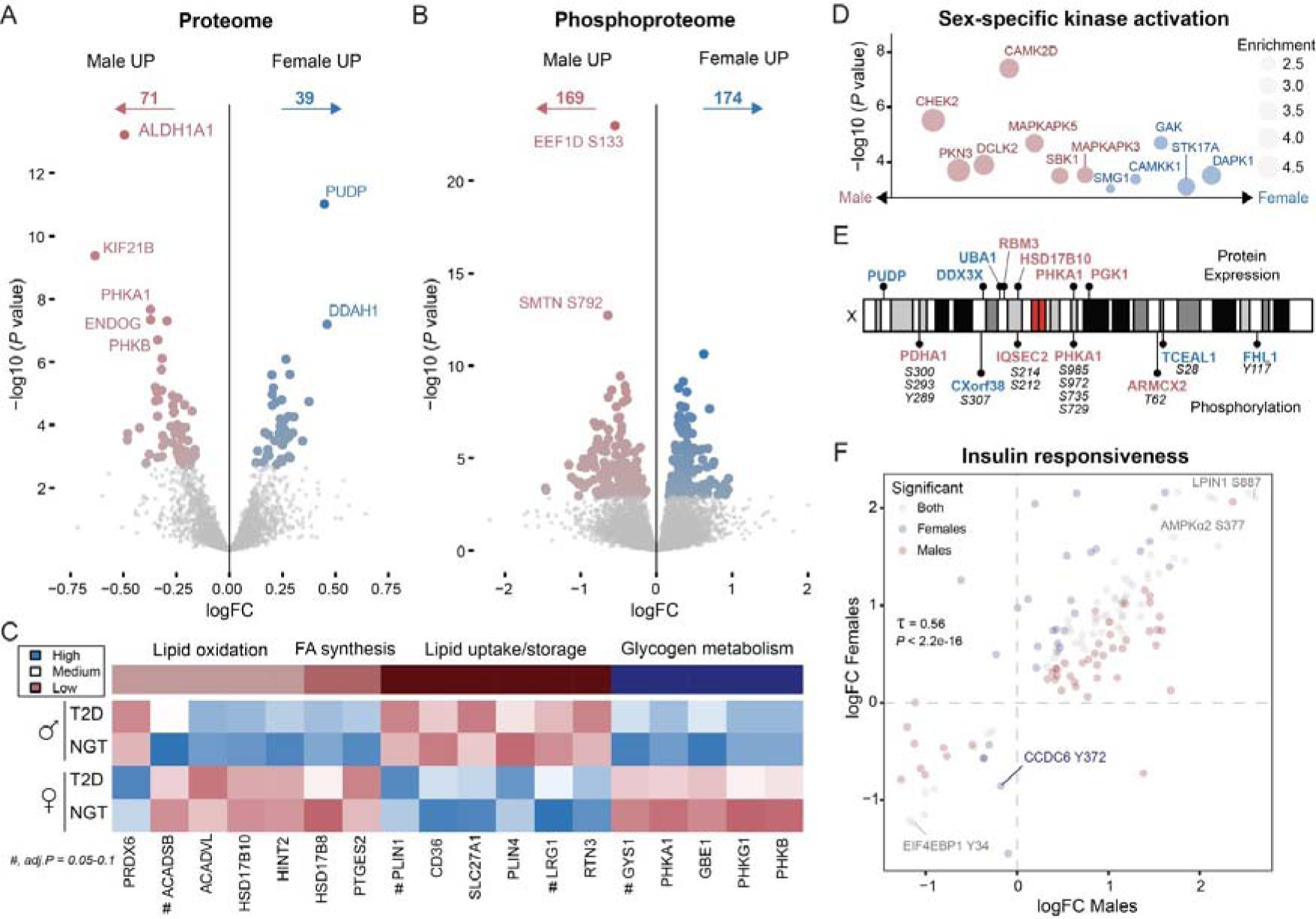
The Sex-Specific Molecular Signature Reveals Shared and Distinct Feature of Metabolism. Volcano plot of differentially expressed proteins (A) and phosphosites (B) between males and females (Limma, main effect < 5% FDR). Significantly regulated proteins related to substrate metabolism including lipid oxidation, fatty acid (FA) synthesis, lipid uptake/storage and glycogen metabolism. # indicates a protein with an FDR value between 5-10% (C). Kinase activity prediction in males and females. Enrichment was performed on all significantly regulated phosphosite (Female vs Male) and the whole phosphoproteome was used as background (D). X-chromosome encoded proteins and phosphoproteins differentially expressed between males (red) and females (blue) (E). Insulin responsiveness in males and females. Kendall’s rank correlation was applied on male, female and shared significant phosphosites regulated by insulin stimulation (F). All data presented in Figure 5 are from the discovery cohort.

A small proportion of the sex-specific proteins and phosphoproteins are located on the X-chromosome (Figure 5E), with some reported to escape female X-chromosome inactivation^50^. Intriguingly, several X-chromosome encoded proteins were higher in skeletal muscle from males, suggesting complete X-chromosome inactivation in females. For instance, HSD17B10 had a higher expression in skeletal muscle from males. HSD17B10 is a key regulator of 17-β-estradiol and androgen metabolism, likely modulating the potency and balance of sex hormone action in skeletal muscle^51,52^. Only seven of the 110 differentially expressed proteins were X-chromosome encoded, suggesting that sex differences largely depend on autosomal and hormonal regulation. This finding is consistent with the observation that few sex-chromosome expressed proteins modulate autosomal gene expression^53^.

Given the substantial sex-specific differences in proteins and phosphosites regulating metabolism, we investigated whether biological sex modulates insulin signaling or the insulin resistant signature in skeletal muscle. For the vast majority of insulin-responsive phosphosites, the response to insulin was comparable between males and females (Figure 5F). Furthermore, sex was not a substantial modifying factor in the association between either proteins or phosphosites with insulin sensitivity (Figure S5I-J). In particular, the abundance of oxidative and glycolytic enzymes was highly coupled to insulin sensitivity in both males and females, despite differences in the abundance of these enzymes between sexes (Figure S5K). Thus, despite major differences in the proteome and phosphoproteome of skeletal muscle, the molecular transducers of insulin action and insulin resistance within skeletal muscle are similar between males and females.

## Discussion

Individuals with type 2 diabetes exhibit impaired whole-body glucose homeostasis, underpinned by reduced insulin-stimulated glucose uptake into skeletal muscle. Despite this knowledge, there are currently no pharmacological therapeutic strategies to specifically target skeletal muscle insulin sensitivity. Furthermore, the detailed molecular atlas explaining insulin action in skeletal muscle, and how this landscape alters during the development of insulin resistance, has been elusive. Resolution of this biology may offer inroads into diabetes treatment strategies, but this has been a challenge owing both to technological limitations and the inherent heterogeneity within type 2 diabetes. Here, we present an advanced proteomics workflow that has enabled the analysis of the proteome and phosphoproteome in an unprecedented number of individuals. Because of this innovation, alongside deep clinical phenotyping, we reveal the personalized molecular signatures of insulin resistance and type 2 diabetes. By interrogating these molecular signatures, alongside measures of diabetogenic traits, we deconvolute the signaling mechanisms coupled to aberrant insulin action in skeletal muscle across varying degrees of insulin sensitivity.

By conducting stratified and personalized proteome phenotype associations, distinct patterns were apparent across a continuum of insulin sensitivity. The role of mitochondria in the development of type 2 diabetes remains contentious^54–56^. Our proteome-phenotype associations reveal that mitochondrial protein content is tightly correlated with whole-body insulin sensitivity, but this signature is not a distinct feature of type 2 diabetes *per se*. Conversely, the abundance of glycolytic enzymes is negatively correlated with insulin sensitivity. Collectively, these data highlight how the molecular landscape of skeletal muscle can determine type 2 diabetes pathophysiology independently of disease diagnosis. By taking direct measures of insulin sensitivity into account, our proteomic analysis has unmasked associations, which are of clinical and therapeutic relevance.

We identified several phosphosites associated with insulin resistance in the fasted state. This was an unexpected discovery because the field has largely focused on the response of target tissues to insulin. Thus, the systemic milieu also appears to provide an environment that triggers changes in the phosphoproteome between diagnosis groups. Hormones and metabolites associated with the diabetogenic state, even under fasting conditions, may provide a permissive stimulus that affects the phosphoproteome and predicts insulin sensitivity. Notably, an unbiased kinase prediction analysis for these phosphosites revealed hyperactivation of the JNK-p38 pathway in insulin-resistant muscle. This phosphoproteomic signature may reflect an altered immunological tone around the skeletal muscle in people with insulin resistance^57^. Strikingly, the phosphosite with the strongest association with insulin sensitivity was on AMPKγ3 S65, a site uniquely found in *homo sapiens*. AMPKγ3 is exclusively expressed in skeletal muscle and is crucial in regulating glycogen metabolism, as well as post-exercise glucose uptake and insulin sensitivity in humans^58–60^. We speculate that S65 could serve as a human-specific rheostat for the regulation of insulin sensitivity. Our data suggest MAPKAPK2 as an upstream kinase for AMPKγ3 S65. Thus, the p38/JNK-MAPKAPK2-AMPKγ3 axis may be a promising therapeutic avenue to improve skeletal muscle insulin sensitivity. Given that AMPKγ3 is muscle-specific, the S65 site could be a potential therapeutic target for the development of skeletal muscle insulin sensitizers, reducing the risk of adverse side-effects in other tissues.

While the fasting phosphoproteomic signature was highly associated with whole body insulin sensitivity, a key feature of type 2 diabetes is an impaired insulin-response within skeletal muscle^8,61^. Indeed, numerous phosphosite-phenotype associations were only apparent in the insulin-stimulated state. For example, several mTORC1 substrates were strongly associated with insulin sensitivity under insulin stimulation, highlighting a node of insulin resistance in type 2 diabetes. Conversely, considerable insulin-stimulated phospho-signaling was apparent even in individuals with the lowest insulin sensitivity. For instance, insulin stimulated the phosphorylation of substrates of AKT, a canonical insulin signaling node^62^, to a comparable degree irrespective of whole-body insulin sensitivity (e.g., GSK3β S9 and TBC1D4 T642). While AKT signaling has been highlighted as essential for GLUT4 translocation^63–66^ and found to be impaired in insulin resistant skeletal muscle^13,67^, this is not a consistent finding^68,69^. Our data indicate that insulin-stimulated AKT activity is preserved in states of insulin resistance or alternatively, a compensatory action of unknown kinases or phosphatases may modulate the phosphorylation of AKT substrates. These observations underscore the emerging appreciation that there is selective insulin resistance along some, but not all signaling nodes^70–72^, warranting a shift in our understanding of dysregulated signaling pathways during insulin resistance. This selective insulin resistance further reinforces the need for comprehensive studies of insulin signaling *in vivo* across diverse states of insulin sensitivity.

Finally, our analysis revealed striking differences in the proteome and phosphoproteome between males and females. Large sex-specific distinctions in the proteome were apparent, with females exhibiting a greater reliance on lipid metabolism, in contrast to males who showed a higher propensity for glucose metabolism, consistent with previous research^73^. Surprisingly, few regulated proteins/phosphoproteins were encoded on the X-chromosome, suggesting a more substantial role for hormonal, whole-body, and autosomal gene regulation in shaping the skeletal muscle molecular landscape. Indeed, cell-autonomous sex differences in the phosphoproteome have been noted in cultured human skeletal muscle cells^74^. However, we did not observe any difference in the molecular signaling of insulin action or the molecular drivers of insulin resistance between the sexes. These data indicate that, despite major differences in metabolism, the mechanisms of insulin resistance are similar between males and females.

Through the application of personalized proteomics, our study leverages heterogeneity in type 2 diabetes pathophysiology to disentangle insulin resistance. We illuminate hidden features of skeletal muscle insulin resistance, unapparent using conventional comparative approaches. In doing so we identify critical molecular pathways associated with insulin resistance. Collectively, these data highlight the need to consider heterogeneity within disease classifications and advocate for a precision medicine approach to type 2 diabetes care.

## Methods

### Human clinical studies

#### Discovery cohort

Individuals with normal glucose tolerance (NGT), as determined by oral glucose tolerance test (OGTT), or type 2 diabetes (T2D), were included in the study. Clinical and transcriptomic data from a subset of males from this cohort has previously been studied^75^. All participants had declared written informed consent, and the study protocol was approved by the regional ethics board (Stockholm, Sweden). Participants were also instructed to avoid physical activity for 48 hours before the experimental day. On the morning of the study, following an overnight fast, anthropometry measures and blood were collected for clinical biochemistry. A biopsy was taken at rest from the *vastus lateralis* muscle using a Weil-Blakesley conchotome instrument (Agnthos, Sweden) under local anesthesia (10 mg/ml; mepivacaine hydrochloride, AstraZeneca, Cambridge, UK). Four-five minutes after resting in a supine position, participants underwent a hyperinsulinemic-euglycemic clamp. Initially, participants received a bolus of insulin (1600 mU/m^2^ body surface area) followed by a constant intravenous insulin infusion at 40 mU/m^2^/min for 2 hours. Simultaneously, an intravenous infusion of glucose was given and adjusted to maintain euglycemia (4.5-5.5 mM). Another muscle biopsy was collected 30 minutes into the clamp. The last 60 minutes of the clamp was considered a steady-state of the glucose infusion rate (GIR) and used to calculate whole-body insulin sensitivity (M-value).

#### Validation cohort

The validation cohort consisted of 34 individuals diagnosed with type 2 diabetes from the “IDA” study and 12 matched subjects exhibiting normal glucose tolerance (NGT), all paired according to sex, body mass index (BMI), age, and smoking status as reported earlier^23^. Prior to their involvement, informed consent was secured from each participant. The ethical oversight for this study was provided by the Regional Scientific Ethical Committees for Southern Denmark, following the principles outlined in the Helsinki Declaration (Project-ID: S-20120186). Preliminary health screening, including blood tests and electrocardiograms (ECG), confirmed the normal health status of all participants. The hyperinsulinemic-euglycemic clamp was performed in the morning after an overnight fast. Participants were instructed to refrain from physical activity for 48 hours before the hyperinsulinemic-euglycemic clamp. For all participants, any glucose, lipid, and blood pressure medications were withdrawn one week in advance of the clamp procedure. The clamp was initiated with a 2-hour basal period involving the infusion of a primed, constant quantity of [3-^3^H]-tritiated glucose to achieve tracer equilibrium. Subsequently, a 4-hour period of insulin stimulation started, employing an insulin infusion rate of 40 mU/m^2^/min. Glucose was infused to maintain euglycemia (5.0-5.5 mmol/l). Muscle biopsies were sampled from the *vastus lateralis* muscle both prior and at the end of the insulin infusion by a modified Bergström needle technique under local anesthesia (lidocaine)^76^.

#### Biopsies

Visible fat, connective tissue, and blood vessels were removed from the muscle biopsies immediately after sampling and frozen in liquid nitrogen. All samples were stored at −80°C until proteomic analysis.

#### Cell culture

Human skeletal muscle primary cells were obtained from ATCC (PCS-950-010) and grown in growth media (ATCC PCS-500-030 supplemented with ATCC PCS-950-040) supplemented with 1% penicillin and streptomycin until 80% confluent. Cells were split into matrix-gel coated 6-well plates. Differentiation was initiated at 90% confluence in differentiation media (ATCC PCS-950-050). At day 6 of differentiation, cells were serum-starved for 4 hours before treated with DMSO or 1µM CC-99677 for 1 hour.

#### Immunoprecipitation

Two days after seeding HEK293T cells in a 6-well plate, they were transfected with 1800 ng of plasmid containing either N- or C-terminal tagged GFP-AMPKγ3 using TransITx2 transfection reagent (Mirus Bio, cat no. MIR 6000) and then lysed 48 hours later (lysis buffer – 50 mM Tris, 1% Triton X-100, 0.27 M sucrose, 1 mM EDTA, 1 mM EGTA, 20 mM glycerol-2-phosphate disodium, 50 mM sodium fluoride, 5 mM tetrasodium pyrophosphate + protease inhibitor cocktail, pH 7.4). GFP tagged AMPKγ3 was then immunoprecipitated by incubating 400 µg of protein with 1 µg of anti-GFP antibody (Chromotek, cat no. 3H9) and 15 µl of protein G sepharose beads (Cytiva, cat no. 17061802) overnight at 4°C on an over-end rotator. The next day the beads were washed two times in lysis buffer followed by three times in TBS. Proteins were eluted and digested in 2 M urea, 50 mM Tris pH = 8.5, 1 m M DTT with 0.5 µg trypsin for 1 hour at 37°C. The supernatant was collected into a new tube and alkylated in 5 mM Iodo-acetamide (IAA), and digestion continued overnight. The next day, digestion was stopped by bringing samples to 1% TFA and 1/5^th^ of eluate loaded onto equilibrated evotips.

### Sample preparation

#### Discovery cohort sample processing

Powdered muscle biopsies were lysed in 4% sodium dodecyl sulfate (SDS), 100 mM Tris pH = 8.5 with an Ultra Turrax homogenizer (IKA) for 2x 10s on ice. Homogenate was immediately boiled at 95°C for 5 minutes at 800 revolutions per minute (RPM). Samples were then sonicated for 30 s (1 s on/off, 50% amplitude) with a tip-probe sonicator before centrifugation for 10 minutes at 16,000g. The supernatant was collected, and protein concentration was determined by the DC assay (Thermo Fischer). Protein lysate was then reduced with 10 mM pH-neutral Tris(2-carboxyethyl)phosphine hydrochloride (TCEP) and alkylated with 40 mM Chloroacetamide (CAA) at 40°C for 5 minutes. Protein aggregation capture was performed on 300 µg protein of SDS lysate per sample in a 2 mL deep 96-well plate on a KingFisher Flex robot (Thermo)^77,78^. Briefly, plate 1 (sample plate) consisted of hydroxyl beads (1:4 protein:bead ratio) (MagResyn, Resyn Biosciences), lysate, and 70% acetonitrile (ACN). Plate 2+3 (wash 1-2) consisted of 100% ACN. Plate 4 (digestion plate) consisted of trypsin (1:100 enzyme: protein) + lysC (1:500 enzyme: protein) in 100 mM Tris pH = 8.5. Plate 1 was mixed for 1 minute, followed by a 10-minute pause for proteins to aggregate to beads. Then, beads were washed 2 times for 2.5 minutes with gentle mixing before being transferred to a digestion plate for overnight digestion at 37°C with medium mixing. The following day, samples were eluted in 1% TFA in two rounds of 10-minute medium mixing. Samples were transferred to 1.5 mL tubes and centrifuged for 10 minutes at 20,000g. Peptides were desalted on equilibrated 50 mg C18 cartridges (Sep-Pak, Waters) and eluted into 1.5 mL tubes. An aliquot for proteomics was transferred to a 96-well plate, dried completely in a speed vacuum centrifuge, and resuspended in 5% ACN, 0.1% TFA. The peptide concentration was measured on a nanodrop and 200 ng was loaded onto equilibrated evotips.

For analysis of the human skeletal muscle myotube phosphoproteome, cells were lysed in 2% SDS, 50 mM Tris pH=8.5 and immediately boiled at 95 for 5 minutes. The lysate was then sonicated (bioruptor) with 10 cycles of 30 s on/off and followed by centrifugation (20.000g, 10 minutes). Protein lysate (75 µg) was then added to kingfisher 2 mL deep-well plate, reduced, alkylated, and digested via the protein aggregation capture protocol as described above.

#### Validation cohort sample processing

SDS protein lysate was obtained as above and precipitated overnight at −20°C with four times volume of acetone. The next day, protein pellet was washed and resuspended in 4% sodium deoxycholate (SDC), 100 mM Tris pH = 8.5 with 15 minutes of 30 s on/off sonication. Reduction and alkylation were performed with 10 mM pH-neutral TCEP and 40 mM CAA and incubated for 5 minutes at 40°C. Protein was digested overnight at 37°C with 1:100 and 1:500 trypsin and lysC, respectively. The following day, digestion was stopped by bringing samples to 1% TFA. SDC precipitate was cleared by centrifugation (20,000g, 15 minutes), and the supernatant was desalted on equilibrated 50 mg cartridges (Sep-Pak, Waters). The elution was done in 50% acetonitrile and concentrated with a speed vacuum drier. The peptide concentration was measured on a nanodrop. 100 µg of peptide was used as input for phosphopeptide enrichment as described below.

#### Phosphopeptide enrichment

Phosphopeptide enrichment was performed on KingFisher flex robot^79^. Peptide (50 µg) was concentrated and loaded into IMAC loading buffer (80% ACN, 5% TFA, 0.1 M glycolic acid) in a deep 96-well plate (sample plate). An equal mixture (15 µL each) of Ti-IMAC-HP and Zr-IMAC-HP beads (MagResyn, Resyn Biosciences) were washed and equilibrated in IMAC loading buffer. The tip plate was located in position #1. The IMAC-bead plate was in position #2. The IMAC-bead washing solution (loading buffer) was placed in position #3, where the sample plate was in position #4. The three washing plates (wash 1: loading buffer, wash 2: 80% ACN, 1% TFA and wash 3: 10% ACN, 0.2% TFA) were placed in positions #5-7, respectively. The Elution plate consisting of 1% NH4OH (freshly prepared) was placed in position #8. The protocol goes as follows: Beads were washed for 5 minutes in IMAC loading (medium speed) buffer followed by 20 minutes of mixing (medium speed) in the sample plate. Beads were then washed once in each washing buffer for 2 minutes (medium speed), followed by 10 minutes of elution in 1% NH4OH. Eluted peptides were then acidified by TFA and loaded directly onto equilibrated evotips.

For phosphoproteomics of primary human muscle cells, resulting tryptic peptides from protein aggregation capture were loaded directly onto an equilibrated C18 96-well plate (Sep-Pak, Waters), washed and eluted in 80% ACN. An aliquot was saved for proteome measurement. Afterwards, phosphopeptide enrichment buffer was added to get the samples to 75% ACN, 5% TFA, and 0.1M GA and enrichment was performed as described above.

#### High-pH fractionation

The library generation (phospho only) was performed by fractionation of 4 µg of phosphopeptides from a pool of all samples via high-pH Reverse-Phase Chromatography utilizing a Kinetex 2.6 µm EVO C18 100Å, 150 x 0.3 mm column (Phenomenex) on an EASY-nLC 1200 system (Thermo) with a flow rate of 1.5 µL/min. The separation was achieved over a 62-minute linear gradient ranging from 3% to 60% of solvent B (10 mM TEAB in 80% acetonitrile) against solvent A (10 mM TEAB in water), with a total run time of 98 minutes, including wash and column equilibration phases. Peptides were eluted into 1.5 µL fractions at a rate of one fraction per 60 seconds resulting in a total of 96 fractions.

Fractions were then concatenated into 48 fractions. Peptides from each fraction were then loaded onto equilibrated Evotips (Evosep).

#### Liquid chromatography mass spectrometry

For the proteome analysis, peptides were separated using a 15-cm column with a 150-μm inner diameter, packed with 1.5 μm C18 beads (Pepsep), on an Evosep ONE HPLC system set to the 30-SPD (30 samples per day) protocol. The column temperature was maintained at 50°C. Eluted peptides were then directed into a timsTOF Pro 2 mass spectrometer (Bruker), using a CaptiveSpray source and a 20-μm emitter, operating in DIA-PASEF mode. For the proteome profiling of non-enriched samples, mass spectrometry data were captured across a 100-1700 m/z spectrum. In the MS/MS stage, each DIA-PASEF cycle lasted 1.8 seconds, spanning an ion mobility spectrum from 1.6 to 0.6 1/K0. Ion mobility calibration was achieved using three specific ions from the Agilent ESI-L Tuning Mix (m/z 622.0289, 922.0097, 1221.9906). The DIA-PASEF utilized a prolonged-gradient method, incorporating 16 DIA-PASEF scans with dual 25 Da windows per ramp, covering a mass range of 400.0-1201.0 Da and an ion mobility range of 1.43-0.60 1/K0. The collision energy decreased linearly from 59 eV at an ion mobility of 1/K0 = 1.3 to 20 eV at 1/K0 = 0.85 Vs cm^2, with both the accumulation and PASEF ramp times fixed at 100 ms.

For the phospho-enriched samples, the DIA-PASEF approach was refined using the py_diAID (Python package for DIA with an automated isolation design)^24^. The mass spectrometry data spanned a 100-1700 m/z range. Within the MS/MS dimension, 20 DIA-PASEF scans covered a 404-1450 m/z spectrum. Each DIA-PASEF scan, with a cycle duration of 2.23 seconds, incorporated two ion mobility windows with variable isolation widths adjusted to precursor densities, setting the mobility spectrum from 1.45 to 0.75 V cm^−2. Collision energy was modulated based on the ion mobility, decreasing from 59 eV at 1/K0 = 1.6 V cm^−2 to 20 eV at 1/K0 = 0.6 V cm^−2, calibrated using three ions from the Agilent ESI Tuning Mix (m/z 622.02, 922.01, and 1221.99), with both accumulation and PASEF ramp times maintained at 100 ms.

#### Data analysis

The phosphoproteome raw files were quantified in Spectronaut v.17.6 in directDIA+ mode against the reviewed FASTA (*Homo sapiens*, 2023 January) with default settings. Phospho(STY) was added as a variable modification and the PTM workflow was enabled with localization probability set to 0. A project-specific phospholibrary was added as a search archive to increase identifications. The phosphosite table was then exported and collapsed in Perseus software, where the site localization probability was set to 0.75^80,81^. The collapsed phosphosite table was then loaded into R studio. The proteome files were quantified in Spectronaut v.18 in directDIA+ with default settings and the output report was directly loaded into R studio.

For the phosphoproteome experiments in human primary myotubes, DIA raw files were processed in directDIA+ mode as above in v.18.6.

For AMPKγ3 pulldown experiments, the raw DDA-files and following MS/MS spectra were quantified and extracted from Fragpipe v20.0^82^. The search was performed with default parameters and with the reviewed human FASTA file (*Homo sapiens*, 2023 January).

#### Bioinformatics analysis

All bioinformatics analyses were performed in the R studio software and graphs were made by using ggplot2. Batch-effects were removed with the limma package and data median normalized^83^. For the proteome data, seven samples were identified as outliers (low IDs) and removed from the data set. Phosphosites and proteins quantified in at least 25% (39 or 37 samples, respectively) of all samples were included in downstream analysis. Inter-subject variation was calculated on raw intensities per protein/phosphosite (fasted and insulin stimulated) across samples as the coefficient of variation. Principal component analysis was performed with the prcomp function (stats package) on the complete quantified protein/phosphosite matrix. Centroid distances of sex, disease and clamp covariates for PC1-15 were calculated, and then z-scored per covariate and plotted as a heatmap.

#### Differentially regulated proteins and phosposites

To identify differentially regulated proteins/phosphosites, we applied the lmFit function with eBayes smoothening (limma package)^83^. Disease state (normal glucose tolerance, type 2 diabetes), clamp (Pre, Post) and sex were added as covariates with blocking for subjects. Phosphosites or proteins with a Benjamini-Hockberg (BH) corrected *P*-value < 0.05 was considered significant. To compare the insulin response between males and females, a separate male/female analysis was performed with disease and clamp added as covariates to the linear model. For analysis of human skeletal muscle myotubes, log2-transformed data was median normalized and HSPB1 S15 and AMPKγ3 S65 site information was extracted. A student’s two-sided t-test was used to test for statistical difference between treatment with DMSO and CC-99677.

#### Correlation analyses

The cor.test function was used to calculate Kendall’s rank correlation between diabetogenic parameters and protein/phosphosite intensities. All analysis were performed across independent samples (i.e. pre and 30-minute time points were analyzed separately). P-values were corrected with the Benjamini-Hochberg method. To compare the proteome and phosphoproteome correlation with insulin sensitivity between males and females, data was split based on sex and separate Kendall’s rank correlations were performed.

#### Summed protein intensities

UseMart and getBM functions were used to retrieve pathway/compartment specific information (biomaRt package)^84^. Proteins related to mitochondria were retrieved with the code GO:0005739^85^. Glycolysis proteins (GO:0006096) were manually filtered for enzymes only part of glycolysis pathway (glucose-pyruvate). Proteins related to the TCA cycle were obtained from the code GO:0006099. Proteins part of mitochondrial complex I-V were retrieved from HUGO Gene Nomenclature Committee (HGNC) database^86^. Oxidative enzymes were defined as TCA and electron-transport chain proteins, whereas glycolytic was defined as enzymes part of the glycolysis pathway. Proportions were calculated based on summed raw intensities.

#### Gene enrichment analyses

GeneSet enrichment analysis was performed with the gseGO/gseKEGG function with default settings (clusterProfiler package)^87^. Benjamini-Hochberg corrected P-values < 0.05 were considered significant. For analysis of AKT and mTOR kinase enrichment, the phosphositeplus (PSP) database of annotated kinase-substrate relationships was used. A Wilcoxon GeneSet enrichment analysis was used to test for kinase insulin sensitivity association (limma package). To assess enrichment of insulin-regulated pathways, a KEGG pathway overrepresentation analysis was performed at the phosphoprotein level. The whole phosphoproteome was used as the background.

#### Clustering analysis

Hierarchical clustering of insulin-regulated phosphosites were performed on z-scored values across groups of insulin sensitivity with the hclust function (stats package). Euclidean distance and average linkage were used as default parameters. The optimal number of clusters was determined using the silhouette analysis. Specifically, the average silhouette width was calculated for cluster solutions ranging from 2 to 10 clusters, and the results were visualized to identify the number of clusters yielding the highest silhouette score.

#### AMPKγ3 structure, gene mapping and FDA-approved drug targets

The structure of AMPKγ3 (Protein ID: Q9UGI9) was obtained from AlphaFold and the S65 residue was manually highlighted^88^. Multiple sequence alignment of N-terminal AMPKγ3 was performed by the online tool “CLUSTALW” (GenomeNet). Gene-chromosome mapping was performed with the biomaRt and karyoploteR packages^84,89^. The list of FDA-approved drug targets was retrieved from the human protein atlas (based on the Drugbank database)^90^.

#### Kinase and regulatory site enrichment

Kinase enrichment analysis was performed as described^91^ from the *in vitro* substrate-Ser/Thr human kinome screen^92^. In brief, we filtered for Ser/Thr-kinases expressed within skeletal muscle at the transcript level^90^. Peptide sequences of significant phosphosites were used as foreground dataset (5% FDR), where the whole phosphoproteome was used as background dataset. For insulin-resistance associated kinases, only the sites where the parent protein was quantified were used. The Ser/Thr-kinase score of the AMPKγ3 S65 site was also based on the *in vitro* the human kinome screen^92^. Regulatory- and disease associated sites information was obtained from the PhosphositesPlus database^93^. A two-sided Fishers exact test was used to test for overrepresentation of regulatory or disease-associated phosphosites.

## Data availability

The mass spectrometry proteomics data have been deposited to the ProteomeXchange Consortium via the PRIDE^94^ partner repository with the dataset identifier PXD049129.

## Article information

## Acknowledgements

Mass spectrometry analyses were performed by the Proteomics Research Infrastructure (PRI) at the University of Copenhagen (UCPH), supported by the Novo Nordisk Foundation (NNF) (grant agreement number NNF19SA0059305). The authors thank Cecilie B. Lindqvist (Novo Nordisk Foundation Center for Basic Metabolic Research) for skilled technical assistance.

## Author Contributions

Conceptualization, JKL, BS, JRZ, AK, and ASD; Human clinical studies, JVS, DA, JB, DEH, KH, MR, JRZ and AK; Investigation, JKL, BS, and JH; Writing – Original Draft, JKL, BS, and ASD; Writing – Review & Editing, All; Funding Acquisition, KH, JRZ, AK, and ASD; Resources, KH, JZ, AK, and ASD; Supervision, KS, JRZ, BS and ASD.

## Inclusion and Diversity

We support inclusive, diverse, and equitable conduct of research.

---

**Figure S1.**
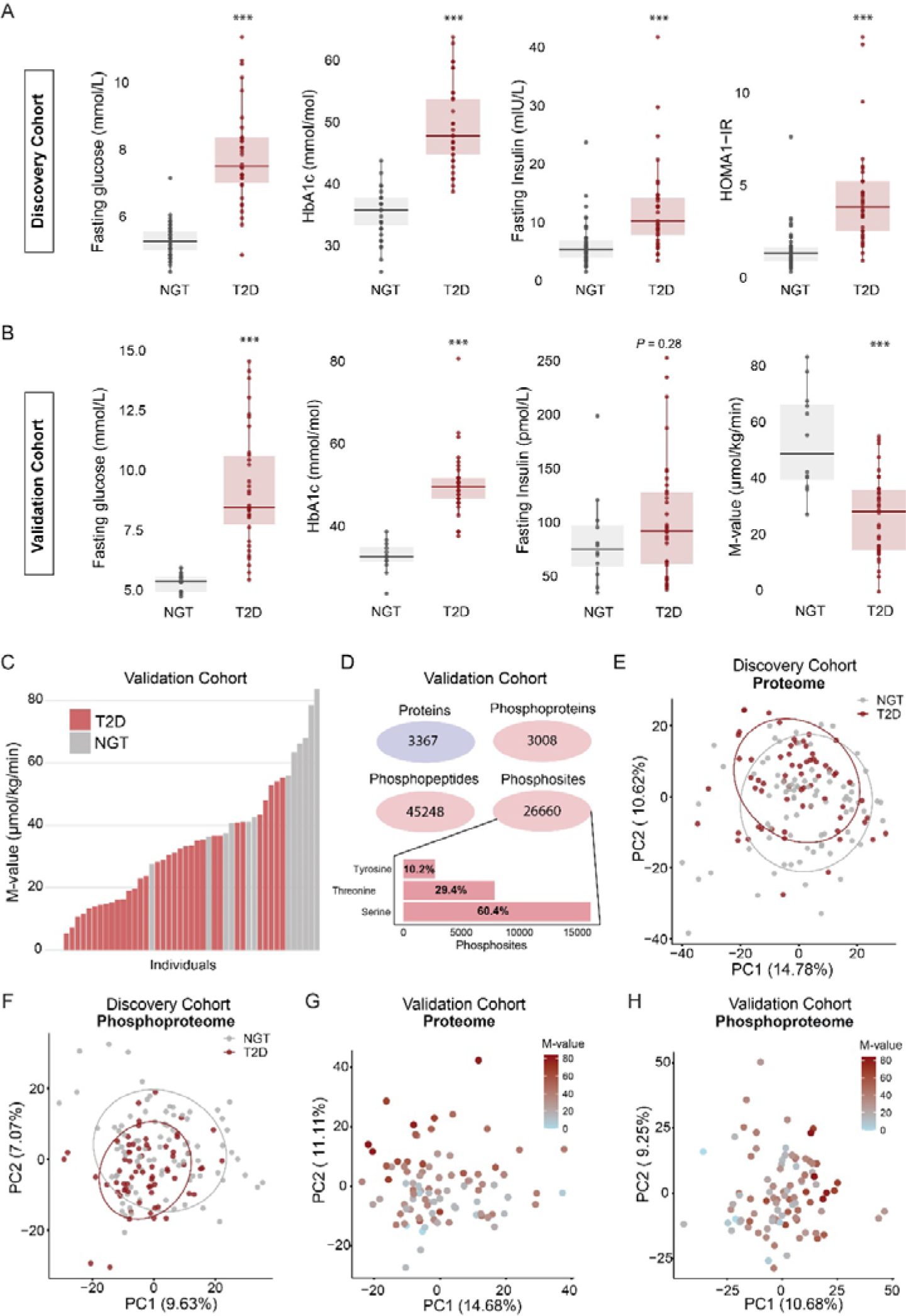
The Proteome and Phosphoproteome of Human Skeletal Muscle are Critical Determinants of Whole-body Insulin Sensitivity. Clinical parameters (fasting glucose, HbA1c, fasting insulin, HOMA1-IR, M-value) for the discovery cohort (A) and validation cohort (B). Insulin sensitivity heterogeneity in the validation cohort (C). Proteomics and phosphoproteomics coverage of proteins, phosphoproteins, phosphopeptides and class 1 (localization probability > 0.75) phosphosites for the validation cohort. Phosphorylation residue distribution between tyrosine, threonine and serine is displayed (D). Principal component analysis proteome (E) and phosphoproteome (F) for the discovery cohort with discrete NGT/T2D group coloring. Principal component analysis of proteome (G) and phosphoproteome (H) for the validation cohort colored by the M-value. A Wilcoxon rank sum test was used in A and B to assess group differences.

**Figure S2.**
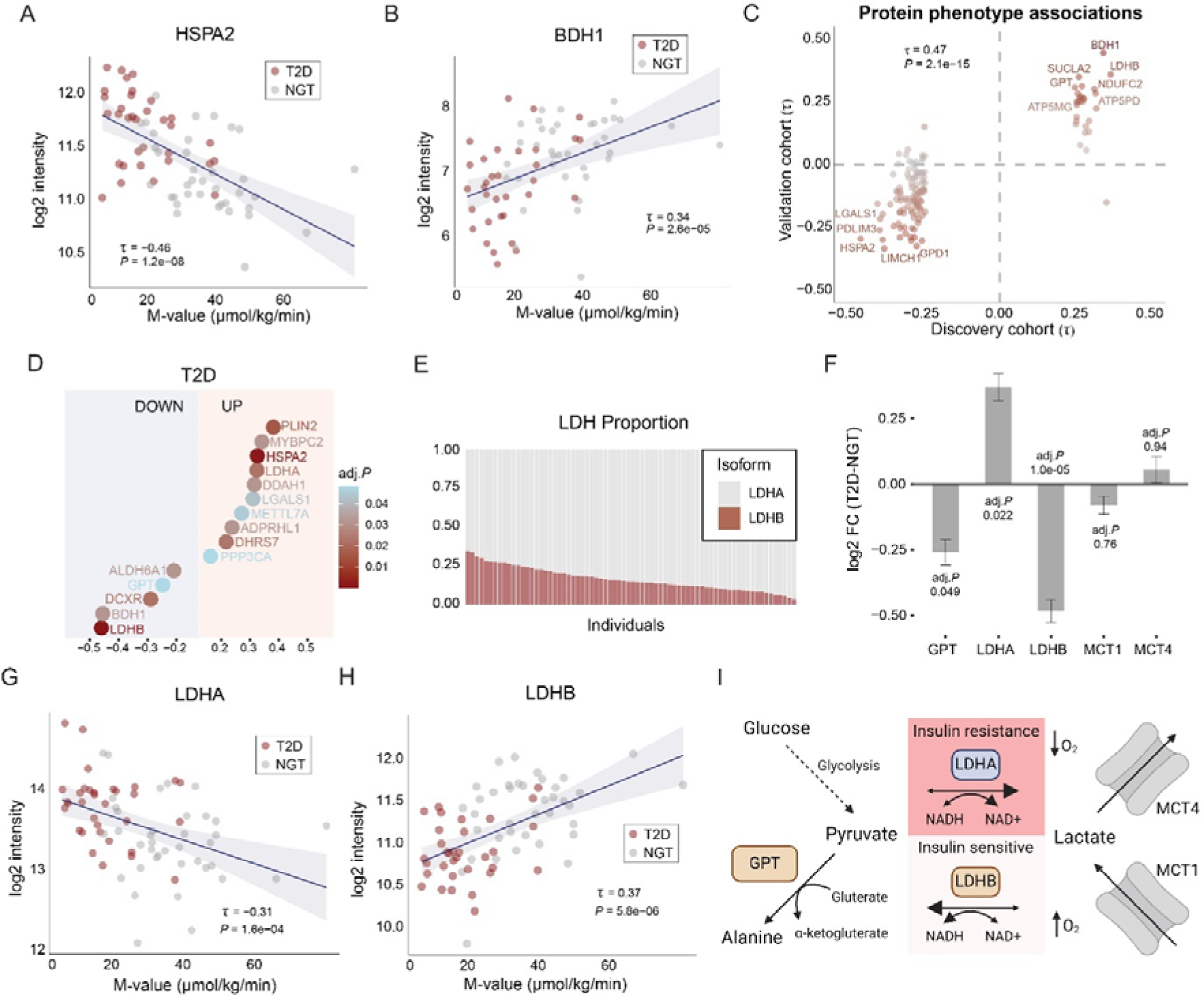
Proteomic Signature of Insulin Sensitive and Resistant Skeletal Muscle. Kendall’s rank correlation of insulin sensitivity (M-value) with baseline protein abundance of HSPA2 (A), BDH1 (B), LDHA (G) and LDHB (H) in the discovery cohort. Correlation of protein phenotype associations between discovery and validation cohort (C). Significantly expressed proteins in a discrete comparison between the T2D and NGT group (Limma main effect, FDR < 5%: discovery cohort) (D). The proportion of Lactate Dehydrogenase (LDH) isoforms across individuals in the discovery cohort (E). Discrete comparison of proteins within pyruvate and lactate metabolism between T2D and NGT group (Limma main effect, FDR < 5%) in the discovery cohort (F). Schematic illustration of pyruvate and lactate metabolism in skeletal muscle (I). Illustration was made in Biorender.

**Figure S3.**
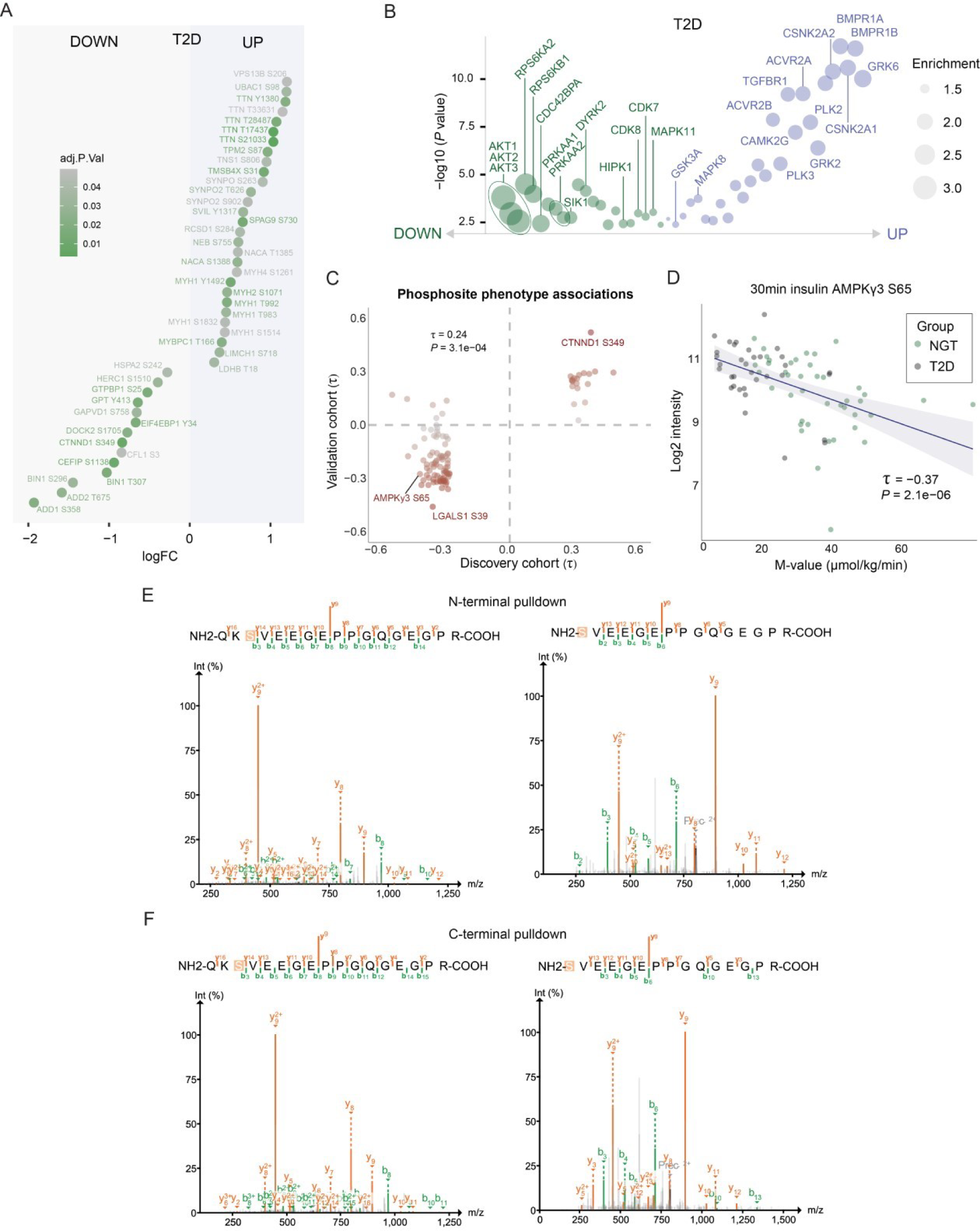
The Fasting Phosphoproteome Landscape is a Critical Determinant of Insulin Sensitivity. Significant regulated phosphosites in a discrete comparison between the T2D and NGT group in the discovery cohort (Limma main effect, FDR < 5%) (A). Kinase activity prediction down (green) and up (blue) in T2D (discovery cohort) (B). Correlation of phosphosite phenotype associations between discovery and validation cohort (C). Kendall’s rank correlation of insulin sensitivity (M value) and AMPKγ3 S65 at 30-minute time point in the discovery cohort (D). Human AMPKγ3-transfected HEK293 cells and subsequent tryptic digestion of AMPKγ3 pull down by N-terminal (E) or C-terminal (F) flag-tag. Peptides were analyzed in data-dependent acquisition mode.

**Figure S4.**
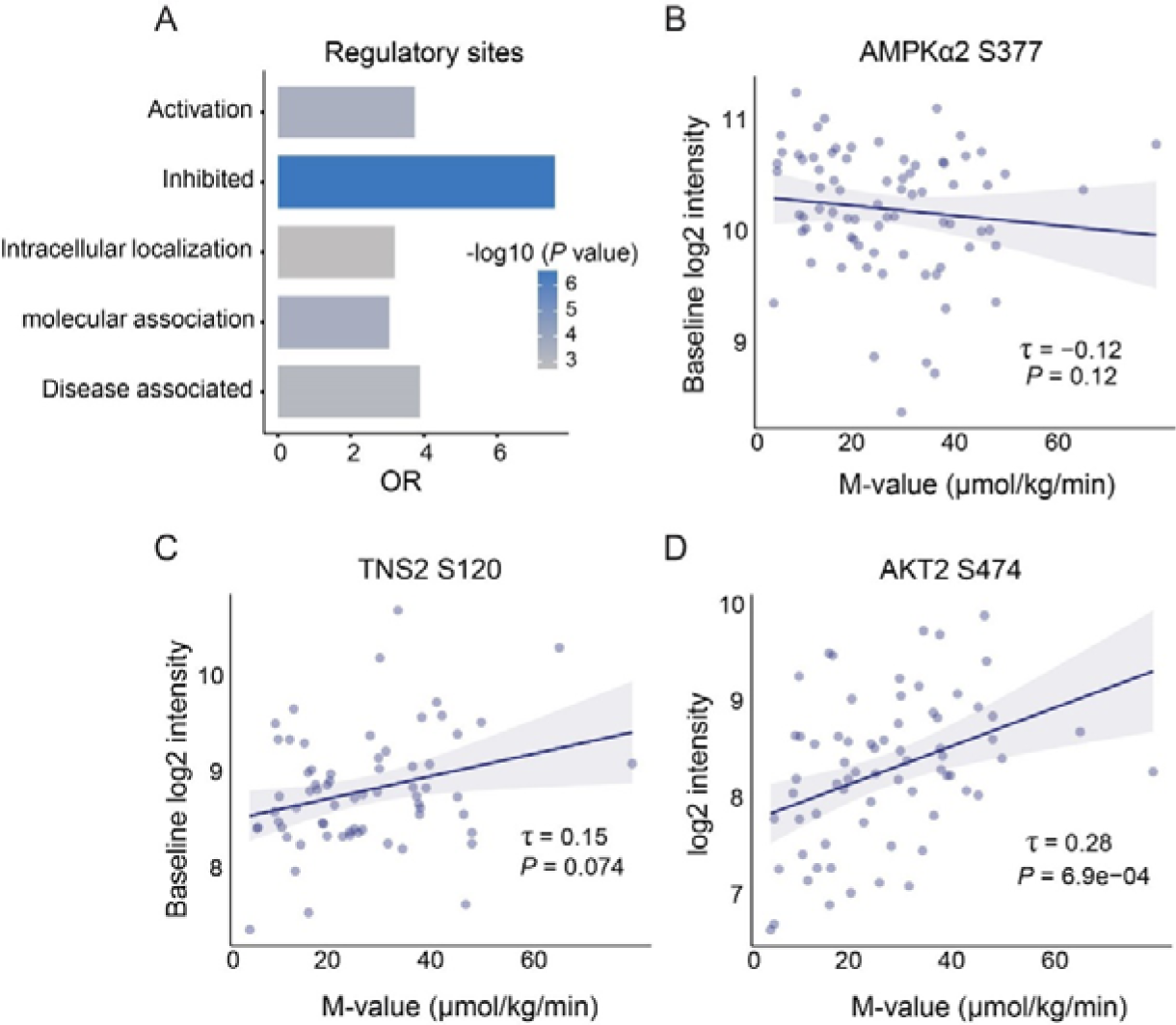
Preserved and Dysregulated Insulin Signaling Across States of Insulin Resistance. (A). PhosphoSitePlus regulatory- or disease-associated sites (two-sided Fishers exact test) (A). Kendall’s rank correlation of insulin sensitivity (M value) and fasting AMPKα2 S377 (B), fasting TNS2 S120 (C) and insulin-stimulated AKT2 S474 log2 intensities (D). OD = Odds Ratio. All data presented in Figure S4 are from the discovery cohort.

**Figure S5.**
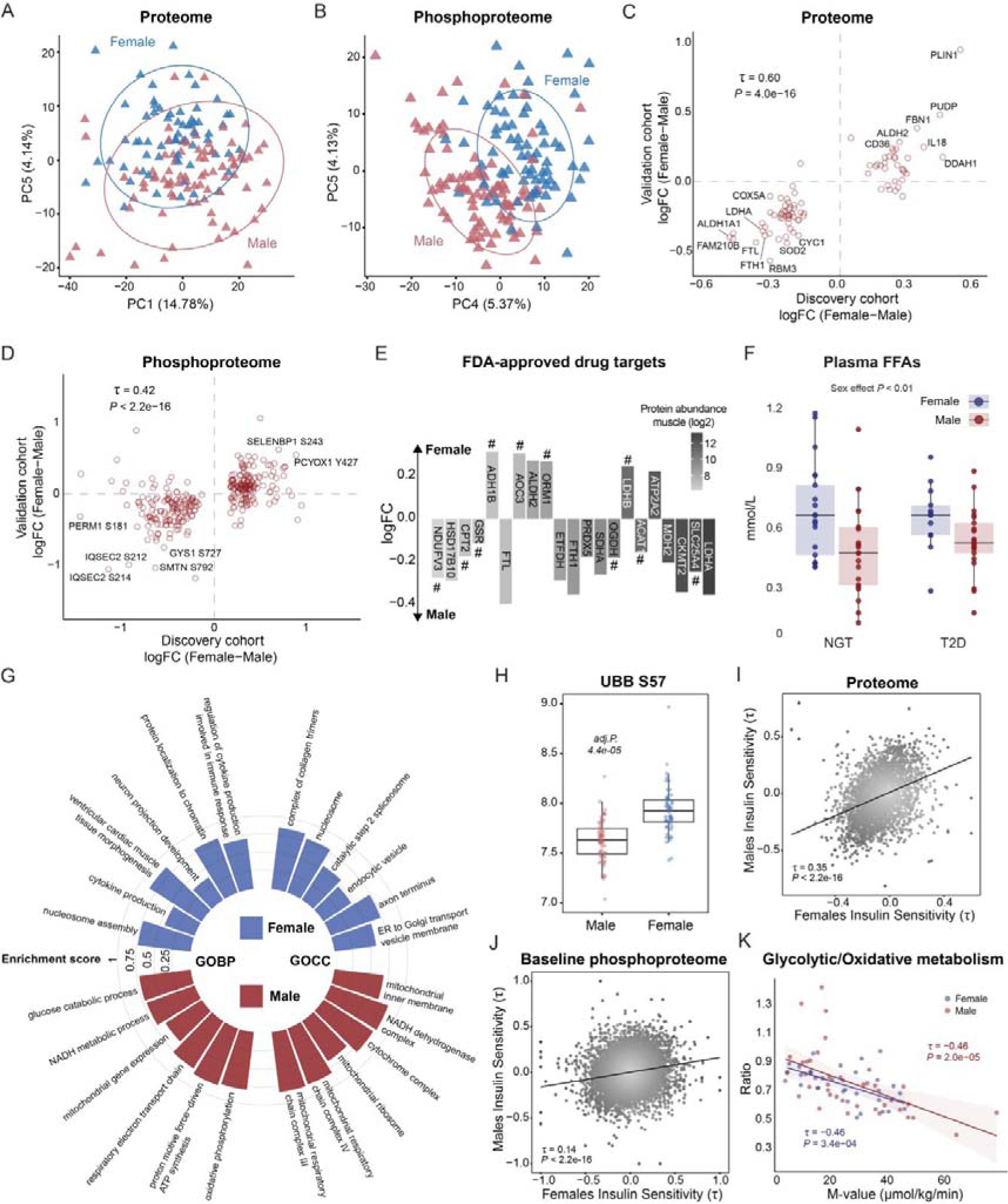
The Sex-specific Molecular Signature Reveals Shared and Distinct Feature of Metabolism. Principal component analysis proteome (A) and phosphoproteome (B) with discrete male/female coloring (discovery cohort). Correlation of female/male logFC differences for proteome (C) and phosphoproteome (D) between discovery and validation cohort. Overlap of sex-specific protein regulation in the discovery cohort and reported FDA-approved drug targets. # indicates proteins that were significant only when combining both the discovery and validation cohort (E). Plasma free fatty acids (FFAs) levels in participants from the discovery cohort. A two-way-ANOVA was used to assess group and sex differences (F). GeneSet enrichment analysis of gene ontology biological processes (GOBP) and cellular components (GOCC) based on logFC differences between females and males in the discovery cohort (G). Boxplots of individual phosphosite log2 intensities in males (red) and females (blue) for Polyubiquitin-B (UBB) S57 in the discovery cohort. The horizontal line indicates the median (H). Insulin sensitivity (M-value) association with baseline proteome (I) and phosphoproteome (J) in males and females from the discovery cohort. The association of glycolytic/oxidative metabolism and insulin sensitivity in males (red) and females (blue) in the discovery cohort. Proteins related to glycolysis and oxidative metabolism (TCA and electron transport chain) were summed and the glycolytic/oxidative ratio was calculated (K). For all correlation analyses, Kendall’s rank correlation was used.

Table S1 Clinical parameters for the discovery and validation cohort

Table S2 Working matrix for phosphoproteome in the discovery cohort

Table S3 Working matrix for proteome in the discovery cohort

Table S4 Working matrix for phosphoproteome in the validation cohort

Table S5 Working matrix for proteome in the validaiton cohort

Table S6 Proteome associations with clinical parameters in the discovery cohort

Table S7 Basal phosphoproteome associations with clinical parameters in the discovery cohort

Table S8 Insulin-stimulated phosphoproteome associations with clinical parameters in the discovery cohort

Table S9 Proteome associations with the M-value in the validation cohort

Table S10 Baseline phosphoproteome associations with the M-value in the validation cohort

